# Time-lapse tryptic digestion: a proteomic approach to improve sequence coverage of extracellular matrix proteins

**DOI:** 10.1101/2025.03.26.645502

**Authors:** Fred Lee, Xinhao Shao, James M Considine, Yu (Tom) Gao, Alexandra Naba

## Abstract

The extracellular matrix (ECM) is a complex and dynamic meshwork of proteins providing structural support to cells. It also provides biochemical signals governing cellular processes, including proliferation, adhesion, and migration. Alterations of ECM structure and/or composition have been linked to many pathological processes, including cancer and fibrosis. Over the past decade, mass-spectrometry-based proteomics has become the state-of-the-art method to profile the protein composition of ECMs. However, existing methods do not fully capture the broad dynamic range of protein abundances in the ECM. They also do not permit to achieve the high coverage needed to gain finer biochemical on ECM proteoforms (*e.g.*, isoforms, post-translational modifications) and topographical information critical to better understand ECM protein functions. Here, we present the development of a time-lapsed proteomic pipeline using limited tryptic proteolysis and sequential release of peptides over time. This experimental pipeline was combined with data-independent acquisition mass spectrometry and the assembly of a custom matrisome spectral library to enhance peptide-to-spectrum matching. This pipeline shows superior protein identification, peptide-to-spectrum matching, and significantly increased sequence coverage against standard ECM proteomic pipelines. Exploiting the spatio-temporal resolution of this method, we further demonstrate how time-resolved 3-dimensional peptide mapping can identify protein regions differentially susceptible to trypsin, which may aid in identifying protein-protein interaction sites.

## INTRODUCTION

The extracellular matrix (ECM) is a remarkably complex scaffold of proteins that plays critical architectural roles in all multicellular organisms, governing cell polarization, organizing cells into tissues, conferring mechanical properties to tissues and organs, and contributing to morphogenesis (1–3). The ECM also conveys biochemical and biomechanical signals tightly controlling all cellular processes, from cell proliferation and survival (4) to migration (5) to differentiation. Alterations of the structure of the ECM scaffold resulting from qualitative or quantitative changes in ECM composition are commonly observed in pathological processes, including skeletal diseases, cardiovascular diseases, fibrosis, and cancer (3, 6, 7).

Over the past decade, bottom-up mass-spectrometry (MS)-based proteomics, a method that infers protein presence based on the detection and identification of the most abundant peptides in a given sample, has helped define the global protein composition of ECMs – or matrisomes – of tissues, organs, or those produced by cells *in vitro* (8, 9). This technology has revealed that the ECM of any given tissue is made of well over 200 distinct proteins and has allowed the identification of ECM protein signatures characteristic of physiological and pathological processes (8, 9).

However, existing proteomic approaches present limitations, especially when using data-dependent acquisition (DDA) mass spectrometry, a modality where only the most abundant precursor ions (peptides) are further fragmented in the second stage of tandem MS and hence identified (8, 10). First, the size and relative abundance of proteins found in the ECM span a very broad dynamic range, from very large and hyper-abundant collagens to small and low-abundance ECM remodeling enzymes or ECM-bound growth factors (8, 11). As such, and despite improvements in instrumentation and off-line enhancements such as sample preparation and protein and peptide fractionation, proteins present in lower abundance are often eclipsed by the more abundant ones (typically fibrillar collagens like COL1A1, COL1A2, COL3A1 in ECM-enriched samples). Additionally, ECM proteins are often highly cross-linked and cannot be easily solubilized, limiting protease accessibility. This is particularly true for structural components of matrisome, *i.e.*, the “core matrisome” that includes collagens, proteoglycans, and other ECM glycoproteins such as fibronectin, fibrillins, tenascins, laminins (11, 12). Last, lower abundance proteins found in the ECM, including ECM-affiliated proteins, ECM regulators, and ECM-bound secreted factors (11), tend to be smaller and thus generated fewer peptides which makes them less likely to be identified. We and others have contributed to the enhancement of proteomic methods, but the focus has remained, until now, to increase the number of proteins identified in a given sample (8, 13). But, like all proteins, ECM proteins exist in multiple proteoforms arising from alternative splicing, single-nucleotide variants, or the presence of post-translational modifications (14), and these proteoforms play key roles in health and disease (13, 15, 16). While proving the presence of a protein in a given sample only requires a few unique peptides, identifying with high confidence proteoforms requires the detection of unique peptide sequences that cover all the variation and modification sites of a given protein, which necessitates higher sequence coverage (13, 17, 18). For reference, the combined average sequence coverage of ECM proteins in MatrisomeDB, a database compiling ECM proteomic datasets (19), is 36.9% with a median sequence coverage of <30%, and of note, the addition of new datasets to MatrisomeDB contributed little to improving overall sequence coverage, as largely similar trypsin-based protocols simply repeat previous observations (19, 20).

Here, we report the development of a “coverage-oriented” approach to achieve deeper matrisome protein sequence coverage. Experimentally, this novel pipeline relies on time-lapsed digestion whereby ECM-enriched samples are incubated in trypsin and released peptides are collected over time. The pipeline also uses data-independent acquisition (DIA) mass spectrometry, a recently developed LC-MS/MS acquisition modality that has gained more traction with improved quantitative reproducibility and broader proteome coverage over the conventional DDA method (21).

Applied to characterize the ECM produced by mouse embryonic fibroblasts in culture, this new pipeline results in a modest but consistent increase in protein identification but, importantly, achieves superior protein sequence coverage compared to standard ECM proteomic protocols. Leveraging the spatio-temporal resolution of this method, we further demonstrate how mapping peptides identified onto the 3-dimensional models of ECM proteins predicted by AlphaFold can identify protein regions differentially susceptible to trypsin that may be the sites of potential protein-protein interactions. Last, since our ability to obtain structures of full-length ECM proteins is limited due to their large size and insolubility, the approach described here can help validate or refine predicted 3D structures of ECM proteins.

## RESULTS

### Design of a time-lapse tryptic digestion pipeline

The process to generate cell-derived matrices (CDMs) by decellularizing the ECM produced by fibroblasts in culture is highly reproducible and thus offers a robust system to test and optimize ECM proteomic workflows. In this study, we used immortalized mouse embryonic fibroblasts (MEFs) to produce CDMs that were partially denatured, deglycosylated, and digested into peptides using either a standard overnight incubation in trypsin or a novel time-lapse digestion protocol including five sampling timepoints: 15, 30, 60, 120, and 240 minutes (**Figure 1A**). Each sample was then analyzed by mass spectrometry using a data-independent acquisition (DIA) modality.

**Figure 1.**
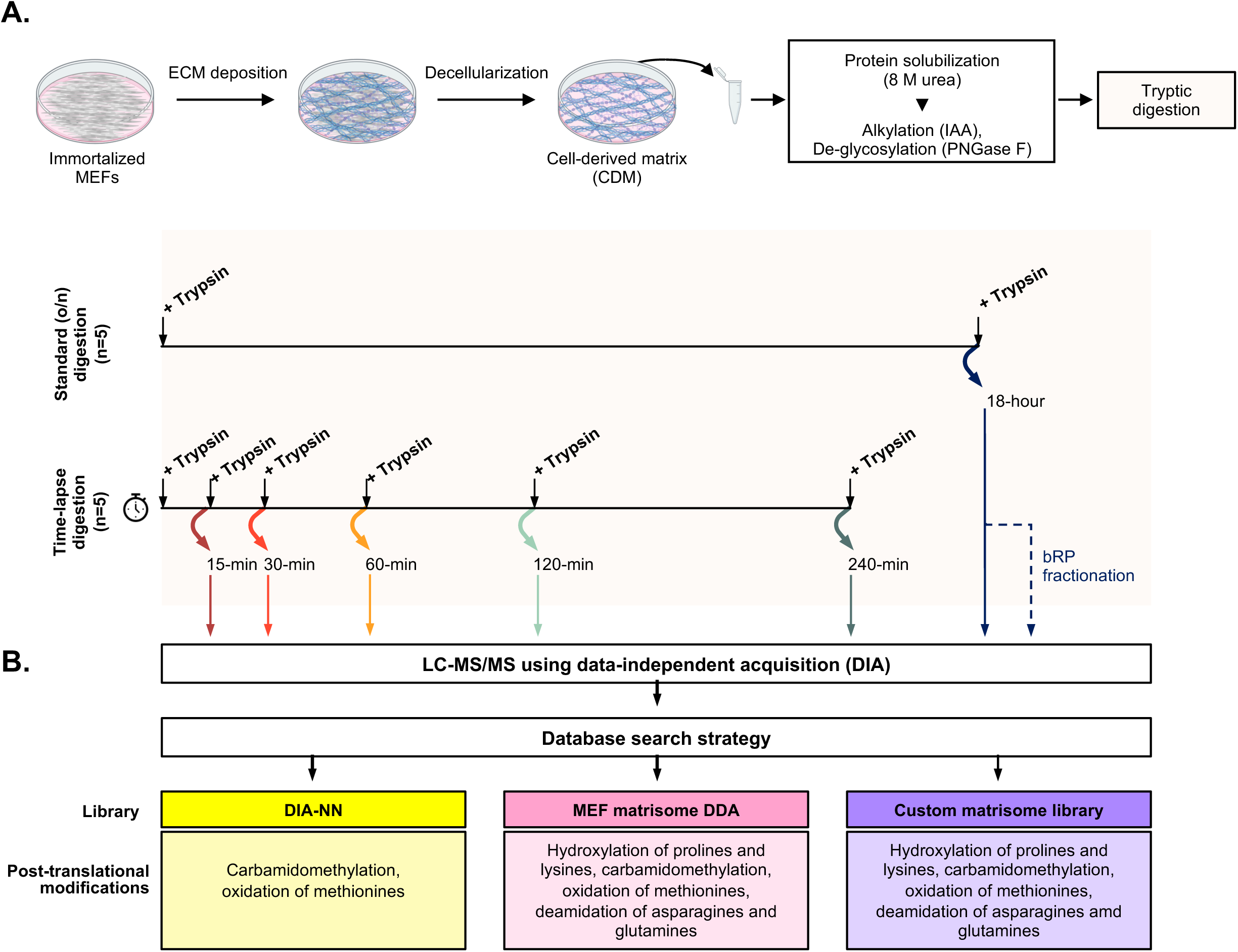
Experimental design. **A.** Schematic representation of the steps to obtain cell-derived matrices (CDMs) from mouse embryonic fibroblasts in culture and process them to generate peptides according to a standard digestion protocol involving an overnight (o/n) incubation with trypsin or a novel time-lapse digestion protocol with sequential tryptic incubations and peptide collections at different timepoints. **B.** Schematic representation of DIA-MSspectra matching process using three approaches relying either on an existing spectral library (DIA-NN), a custom-built spectral library generated from a previous dataset acquired using a data-dependent mode (DDA), or an enhanced library (custom matrisome library). Post-translational modifications allowed in each search are listed.

### Custom spectral library enhances the identification of ECM proteins in DIA datasets

DIA is a modality that has recently emerged and presents certain advantages compared to DDA, including broader proteome coverage and better quantitative reproducibility for known samples (21). In brief, while DDA only selects the most abundant precursor ions (*i.e.*, peptides) for fragmentation, in DIA, all peptides are fragmented and analyzed in a second stage of tandem mass spectrometry. As a result, fragment ion spectra are highly multiplexed and matching spectra to peptide sequences presents some challenges and is highly dependent on the spectral library used (21, 22). Hence, choosing the right spectral library has a significant impact on the output. Interestingly, we noted that, while in DDA mass spectrometry collagen peptides contribute the largest proportion of total precursor ion intensity (“signal”) of any given ECM-enriched sample (8, 9, 19), the proportion of the signal intensity attributable to collagen signals upon DIA, using the DIA-NN spectral library (23) was lesser (∼30%) than the proportion attributable to ECM glycoproteins (**Figure 2A, left panel; red arrow**). One caveat of the predicted library is the inclusion of PTMs, which is not currently well supported by the DIA-NN tool, especially for non-common PTMs, such as proline and lysine hydroxylation. We thus sought to assemble custom spectral libraries and search pipelines (including post-translational modifications unique to ECM proteins such as hydroxylation of prolines and lysines (13, 15, 24, 25)) to ensure capturing matrisome peptides not sufficiently represented in the DIA-NN library (**Figure 1B**). The first custom library was generated using a DDA-MS dataset acquired previously on a pilot time-lapse digestion of samples similar to those analyzed here, namely, cell-derived matrices produced by MEFs in culture (*see also Materials and Methods*). We also assembled a meta-library, hereafter referred to as “custom matrisome library,” composed of the DIA-NN library, the DDA library, and a library derived from a few published datasets on the matrisome of the lung ECM published by Schiller and colleagues (26) (**Supplementary Table S1**).

**Figure 2.**
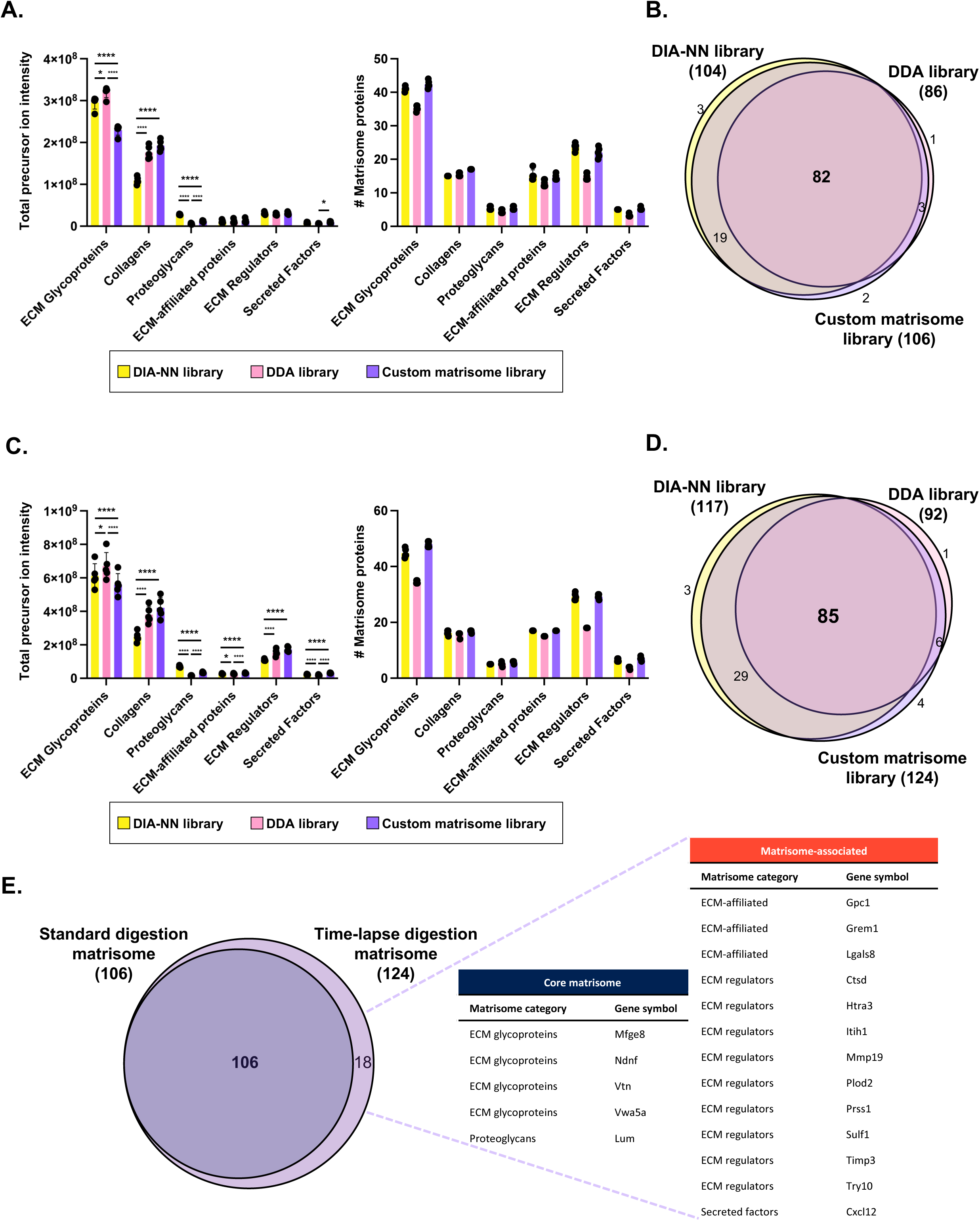
A custom spectral library enhances the identification of matrisome proteins in DIA datasets. **A.** Bar graphs represent the total precursor ion intensities measured (left) and the number of proteins identified (right) by searching the data generated using the standard overnight tryptic digestion pipeline against each of the three spectral libraries built for this study and for each category of matrisome proteins (n=5). Statistical significance is depicted as follows: * p<0.05, **** p<0.0001, the absence of value indicates no statistical significance. **B.** Venn diagram represents the overlap between the matrisome proteins identified in at least three of the five independent biological replicates by searching the data generated using the standard overnight tryptic digestion pipeline against each of the three spectral libraries built for this study. **C.** Bar graphs represent the total precursor ion intensities measured (left) and the number of proteins identified (right) by searching the aggregated data generated using the time-lapse tryptic digestion pipeline against each of the three spectral libraries built for this study and for each category of matrisome proteins (n=5). Statistical significance is depicted as follows: * p<0.05, **** p<0.0001, the absence of value indicates no statistical significance. **D.** Venn diagram represents the overlap between the matrisome proteins identified in at least three of the five biological replicates by searching the aggregated data generated using the time-lapse tryptic digestion pipeline against each of the three spectral libraries built for this study. **E.** Venn diagram represents the overlap between the matrisome proteins identified by searching the data generated through the standard overnight tryptic digestion pipeline (dark purple, left) or the aggregated data generated using the time-lapse tryptic digestion pipeline (little purple, right) against the custom matrisome spectral library. The lists of proteins composing the standard digestion matrisome and the time-lapse digestion matrisome include proteins found in at least three of the five biological replicates.

To determine the impact of spectral library composition on protein identification, we first searched the data collected on peptides generated using a standard overnight tryptic digestion against all three libraries. We found that the DIA-NN library yielded less collagen signal but more proteoglycan signal than the search using the DDA or custom matrisome library (**Figure 2A, left panel** and **Supplementary Table S2**). We also observed that the proportion of the total precursor intensity attributed to ECM glycoproteins was the greatest when searching the dataset against the DDA library (**Figure 2A, left panel** and **Supplementary Tables S2A-C**). Yet, in terms of the number of matrisome proteins identified, using the DIA-NN or the custom matrisome library proved superior to using the DDA library, specifically for the identification of ECM glycoproteins and ECM regulators (**Figure 2A, right panel** and **Supplementary Tables S2A-C**). Since proteomic output presents a certain level of stochasticity, we applied stringent criteria to accept protein identification (protein identification only accepted with at least two unique peptides, strict false-discovery rates at the protein and peptide levels; *see Methods*) and further restricted our analysis to proteins detected in at least three of the five independent biological replicates analyzed. In doing so, we found that an overall larger number of matrisome proteins was identified when the dataset was searched against the DIA-NN or custom matrisome library (104 and 106, respectively) compared to the search against the DDA library (86 proteins), with 82 matrisome proteins identified by all three libraries (**Figure 2B** and **Supplementary Table S2D**).

We next analyzed the dataset acquired on samples prepared using the novel time-lapse tryptic digestion pipeline. We first observed that this pipeline allowed a deeper profiling, since the total precursor ion intensity of matrisome components was twice that obtained with the standard digestion protocol (**Figure 2C, left panel** and **Supplementary Tables S2A-C**). Similarly to what we observed with the standard digestion pipeline, we found that the DIA-NN library yielded less collagen signal, but more proteoglycan signal compared to the search using the DDA or custom matrisome library (**Figure 2C, left panel** and **Supplementary Tables S2A-C**). Interestingly, we also noted that an increasing proportion of the precursor ion signal intensity could be attributed to peptides from ECM regulators at later digestion timepoints (60, 120, and 240 minutes), independently of the spectral library searched (**Supplementary Figure S1**). In terms of matrisome protein identification, the DIA-NN and custom matrisome libraries identified similar numbers of proteins across the different matrisome categories, while the DDA library identified fewer matrisome proteins (**Figure 2C, right panel** and **Supplementary Table S2E**). Comparison of the list of matrisome proteins detected in at least three of the five biological replicates analyzed showed that an overall larger number of matrisome proteins was identified when the dataset was searched against the DIA-NN or custom matrisome library (117 and 124, respectively) compared to the search performed against the DDA library (92 proteins), with 85 matrisome proteins identified by all three libraries (**Figure 2D and Supplementary Tables S2D-E**).

### Time-lapse tryptic digestion moderately increases protein identification

We next compared the list of matrisome proteins detected in at least three of the five biological replicates of the standard or the time-lapse digestion pipelines using the custom matrisome library and found that the time-lapse digestion pipeline identified a larger number of matrisome proteins (124) than the standard pipeline (106) (**Figure 2E**). Five of the 18 additional proteins identified were core matrisome proteins, including the von Willebrand factor A domain containing 5A (Vwa5a), vitronectin (Vtn), and the proteoglycan lumican (Lum), and 13 were matrisome-associated proteins, including enzymes like the matrix metalloproteinase 19 (Mmp19), the procollagen-lysine, 2-oxoglutarate 5-dioxygenase 1 (Plod1, encoding lysyl hydroxylase 1), and the tissue inhibitor of metalloproteinase-3 (Timp3) (**Figure 2E**).

### Time-lapse tryptic digestion increases protein sequence coverage

After evaluating the impact of the two digestion pipelines on matrisome protein identification, we sought to compare the methods in terms of protein sequence coverage, the percentage of a given protein sequence matched (or “covered”) by peptides detected via mass spectrometry. Line graphs presented in **Figure 3A** depict the cumulative sequence coverage achieved through the course of the time-lapse tryptic digestion for each category of matrisome proteins (**Supplementary Table S3A**) and show that, for most matrisome proteins, sequence coverage increases with the addition of peptides released over time. We also observed that the 30-minute timepoint resulted in a sharper increase in sequence coverage across all matrisome categories (Figure 3A).

**Figure 3.**
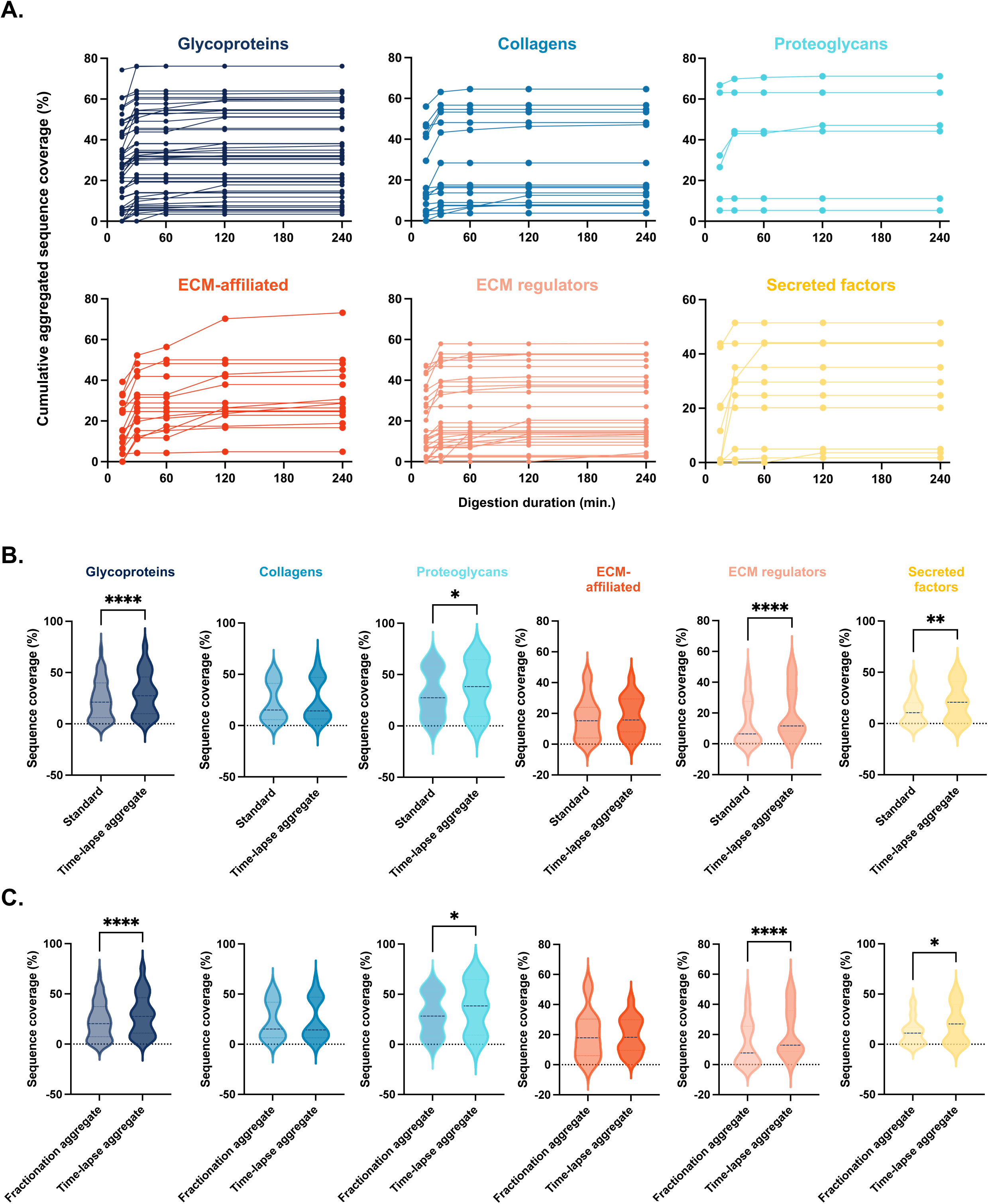
Aggregation of data from different digestion timepoints increases sequence coverage. **A.** Line graphs represent the cumulative average sequence coverage achieved throughout the time-lapse tryptic digestion for each category of matrisome proteins. **B.** Violin plots compare the distribution of the average sequence coverage achieved using the standard tryptic digestion pipeline or the average sequence coverage achieved in aggregate with the time-lapse tryptic digestion pipeline. Statistical analysis was performed using a Wilcoxon signed-rank test, and statistical significance is depicted as follows: * p<0.05, ** p<0.01, **** p<0.0001. **C.** Violin plots compare the distribution of the average sequence coverage achieved in aggregate using basic reversed-phase fractionation of samples generated with the standard tryptic digestion pipeline or the cumulative average sequence coverage achieved with the time-lapse tryptic digestion pipeline. Statistical analysis was performed using a Wilcoxon signed-rank test, and statistical significance is depicted as follows: * p<0.05, **** p<0.0001.

To evaluate whether a particular matrisome category of proteins benefitted more from this approach, we compared the coverage values obtained using the standard overnight digestion or after aggregating coverage data of the time-lapse pipeline (**Figure 3B**). We observed a shift in the distribution of coverage values, particularly for ECM glycoproteins (p < 0.0001), proteoglycans (p < 0.05), ECM regulators (p < 0.0001), and secreted factors (p < 0.01) (**Figure 3B**).

The time-lapse digestion pipeline can be assimilated to a peptide fractionation approach. Since peptide fractionation approaches tend to result in an increased number of proteins identified (9, 27, 28), we next sought to benchmark the time-lapse digestion dataset against a dataset obtained on samples processed through the standard overnight tryptic digestion pipeline and further fractionated using basic reversed-phase fractionation (bRP), a state-of-the-art method for peptide fractionation prior to LC-MS/MS. As expected, bRP fractionation resulted in a greater number of matrisome proteins identified over the standard digestion pipeline and the time-lapse digestion pipeline **(Supplementary Figure S2)**. We next compared the coverage values obtained upon bRP fractionation or after aggregating coverage data of the time-lapse pipeline and observed a statistically significant increase in the distribution of coverage values in the time-lapse dataset, particularly for ECM glycoproteins (p < 0.0001), proteoglycans (p < 0.05), ECM regulators (p < 0.0001), and secreted factors (p < 0.05) (**Figure 3C**).

Upon calculation of cumulative coverage, we found that the highest coverage was achieved for fibronectin (75.6%), while fibronectin’s coverage was 60.8% using the bRP pipeline; this modest increase was nonetheless statistically significant (p=0.024) (**Supplementary Table S3C**). To further identify the proteins that benefitted the most from the increase in coverage achieved by employing the time-lapsed digestion pipeline, we calculated for each protein the ratio of coverage achieved through the aggregation of the data from the different timepoints from the time-lapse digestion pipeline to the coverage attained through standard digestion (**Supplementary Table S3B**) or standard digestion followed by bRP fractionation (**Supplementary Table S3C**). The top five proteins that showed the largest and statistically significant increase in coverage using the time-lapse digestion over the standard protocol were the ECM glycoprotein thrombospondin 4 (Thbs4; 4.5 fold-change increase; p = 0.0002), the matrix metalloproteinase 14 (Mmp14; 4 fold-change increase; p < 0.0001), ABI family member 3 binding protein (Abi3bp; 3.8 fold-change increase; p = 0.0171), fibulin 7 (3.4 fold-change increase; p = 0.0055), and fibroblast growth factor 2 (Fgf2; 3.1 fold-change increase; p < 0.0001) (**Figure 4A, Supplementary Table S3B**). The top five proteins that showed the largest and statistically significant increased coverage using the time-lapse digestion over the bRP fractionation protocol were the ECM regulator procollagen-lysine, 2-oxoglutarate 5-dioxygenase 1 (Plod1; 4.5 fold-change increase; p = 0.0002), Slit3 (3.2 fold-change increase; p = 0.0001), the a1 chain of collagen VIII (Col8a1; 4.5 fold-change increase; p = 0.047), fibroblast growth factor 2 (Fgf2; 2.9 fold-change increase; p < 0.0001), and Mmp14 (2.4 fold-change increase; p < 0.0001) (**Figure 4B, Supplementary Table S3C**).

**Figure 4.**
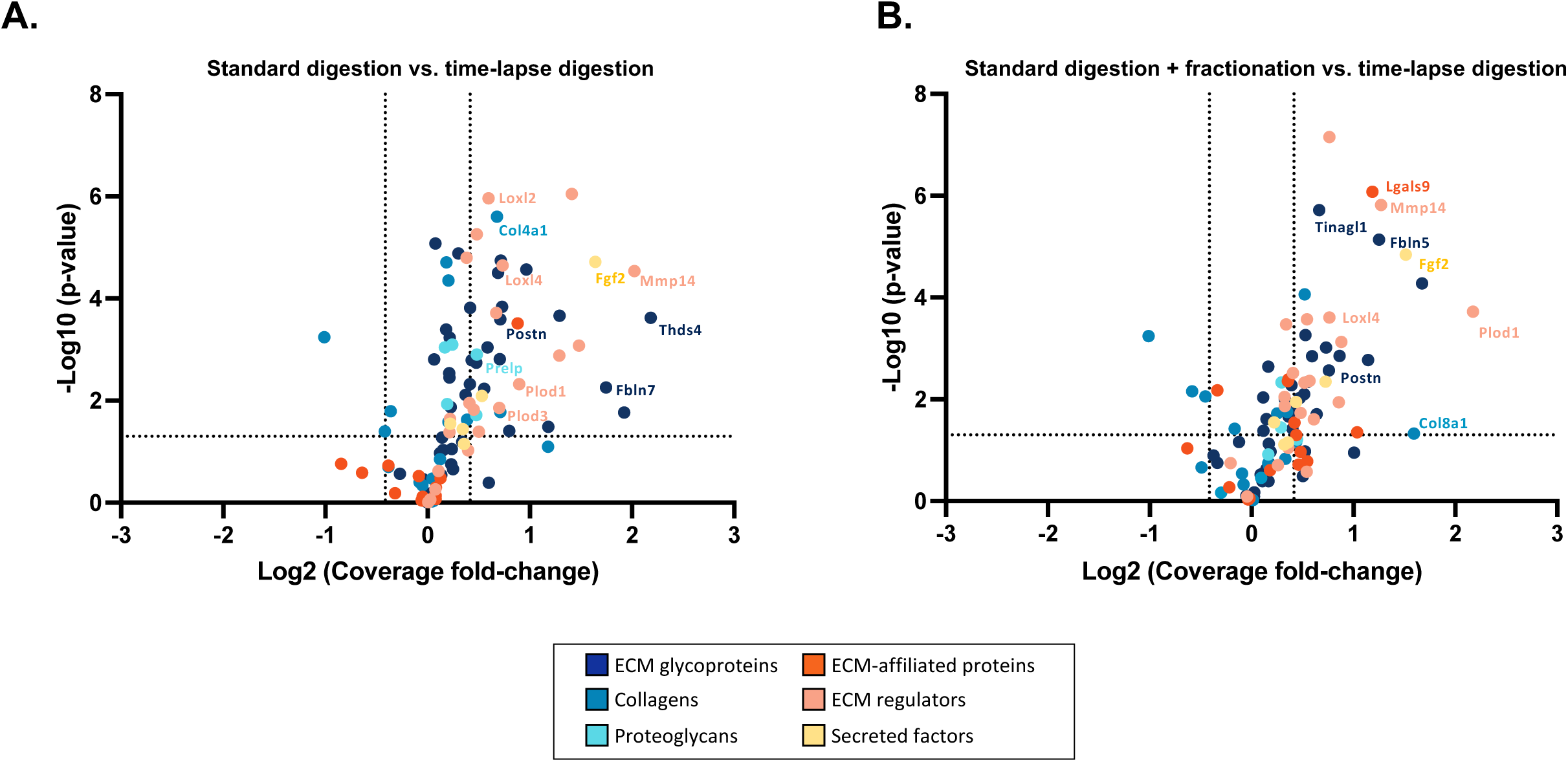
Aggregation of data from different digestion timepoints increases sequence coverage. **A.** Volcano plot represents the increase in sequence coverage attained upon aggregation of the data obtained at each timepoint of the time-lapse tryptic digestion compared to standard overnight tryptic digestion. Dotted lines along the x-axis indicated a 20% increase (right) or decrease (left) in sequence coverage; the dotted line along the y-axis depicts the statistical cut-off set as p<0.05. Proteins above this dotted line are identified with statistically significant changes in coverage in the time-lapse dataset compared to the standard digestion dataset. **B.** Volcano plot represents the increase in sequence coverage attained upon aggregation of the data obtained at each timepoint of the time-lapse tryptic digestion compared to standard overnight tryptic digestion followed by peptide fractionation via basic reversed-phase chromatography (bRP). Dotted lines along the x-axis indicated a 20% increase (right) or decrease (left) in sequence coverage; the dotted line along the y-axis depicts the statistical cut-off set as p<0.05. Proteins above this dotted line are identified with statistically significant changes in coverage in the time-lapse dataset compared to the standard digestion dataset.

In conclusion, while performing time-lapsed tryptic digestion and aggregating data results in only a modest increase in protein identification, it leads to the identification of a significantly larger number of peptides that permits a more accurate estimation of relative protein abundance and achieves a significant increase in protein sequence coverage.

### Sequential release of peptides provides insight into protein accessibility to trypsin and protein conformation

Perhaps the most significant innovation from the time-lapsed digestion protocol presented here is its spatio-temporal resolution. Protein digestibility reflects the surface exposure of a protein and is directly linked to protein folding and protein-protein interactions. Thus, time-lapsed peptide mapping can identify protein regions readily exposed and accessible to trypsin (peptides matching these regions would be released at early digestion timepoints) and regions of proteins hard to access, either because of extensive post-translational modifications, such as O-glycosylations hindering accessibility, or because the engagement of protein-protein interactions not disrupted by the partial denaturation to which samples are subjected. To visualize this, we next thought to map the peptides identified onto 3-dimensional models of ECM protein structures predicted by AlphaFold (29) using an algorithm we recently developed called Sequence Coverage Visualizer (SCV) (30). To illustrate the information that can be leveraged from this visualization, we selected a set of proteins representative of the matrisome, including the glycoprotein periostin (Postn; **Figure 5**), the α1 chain of collagen VIII (Col8a1; **Figure 6**), the proteoglycan prolargin (Prelp; **Figure 7**), and the ECM regulator lysyl oxidase-like 4 (Loxl4; **Figure 8**). In all examples, we provide the theoretical coverage computed via *in-silico* digestion. This allows us to determine whether some proteins are intrinsically resistant to trypsin due to the under-representation of lysines or arginines, the cleavage sites of trypsin. Here, the theoretical coverage ranged from 77.7% for Col8a1 to over 97% for Postn, Prelp, and Loxl4, demonstrating the potential digestibility of the proteins.

**Figure 5.**
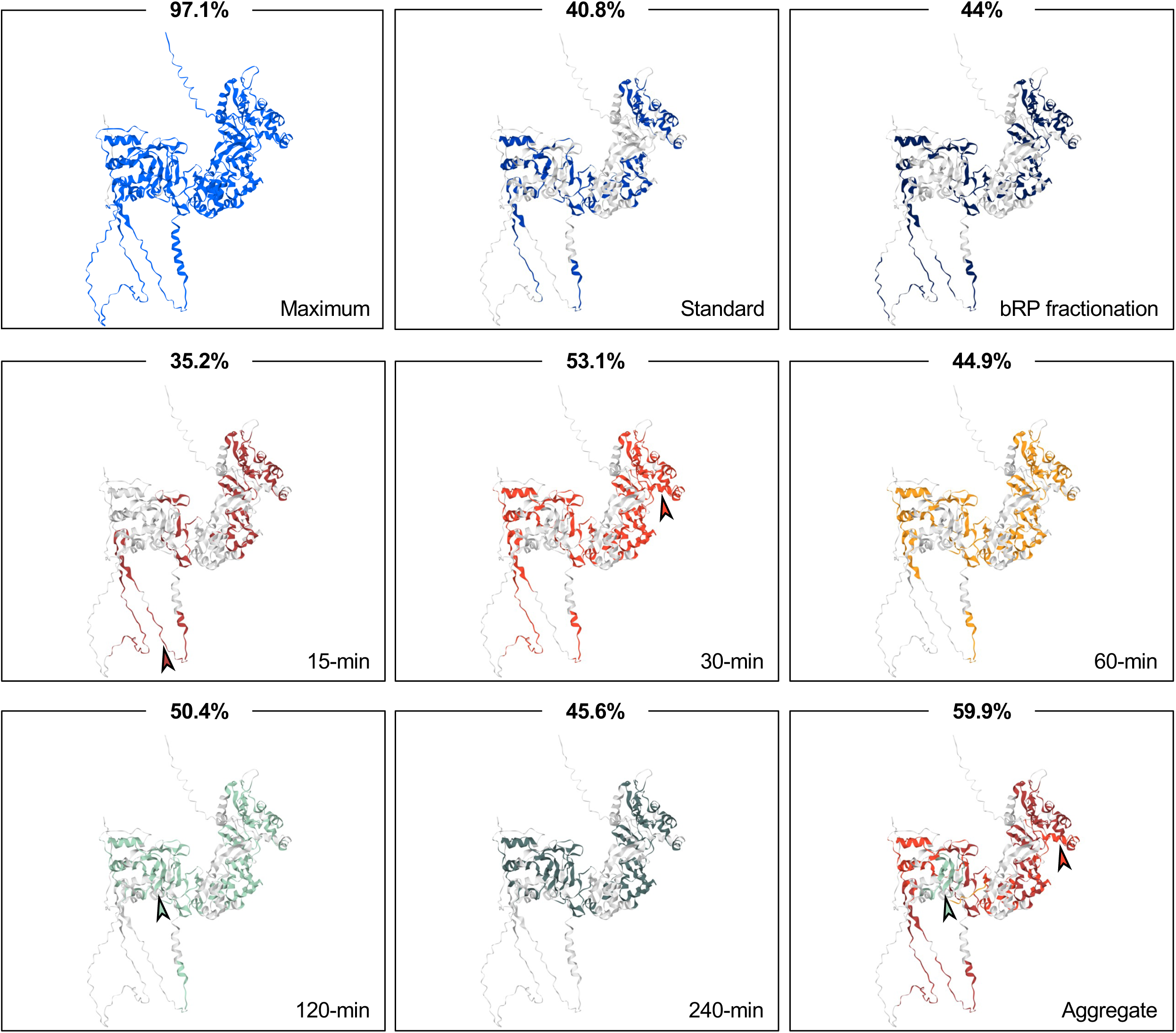
3D visualization of the sequence coverage of the ECM glycoprotein periostin achieved using different experimental pipelines. Panels represent the overlay of peptides detected using the standard tryptic digestion pipeline, the bRP fractionation pipeline, or at different timepoints (15 min: dark red, 30 min: light red, 60 min: gold, 120 min: teal, and 240 min: dark green) of the time-lapse tryptic digestion pipeline, onto the AlphaFold-predicted structure of periostin (UniProt Q62009; https://alphafold.ebi.ac.uk/entry/Q62009). The sequence coverage indicated for each experimental condition was calculated as the proportion of the sequence covered by peptides detected in at least three of the five biological replicates. The “maximum” panel depicts the theoretical sequence coverage predicted via *in-silico* digestion. Peptides are color-coded in the “Aggregate” panel based on the first timepoint at which they were detected. Arrowheads point to peptides uniquely contributed by a given tryptic digestion timepoint.

**Figure 6.**
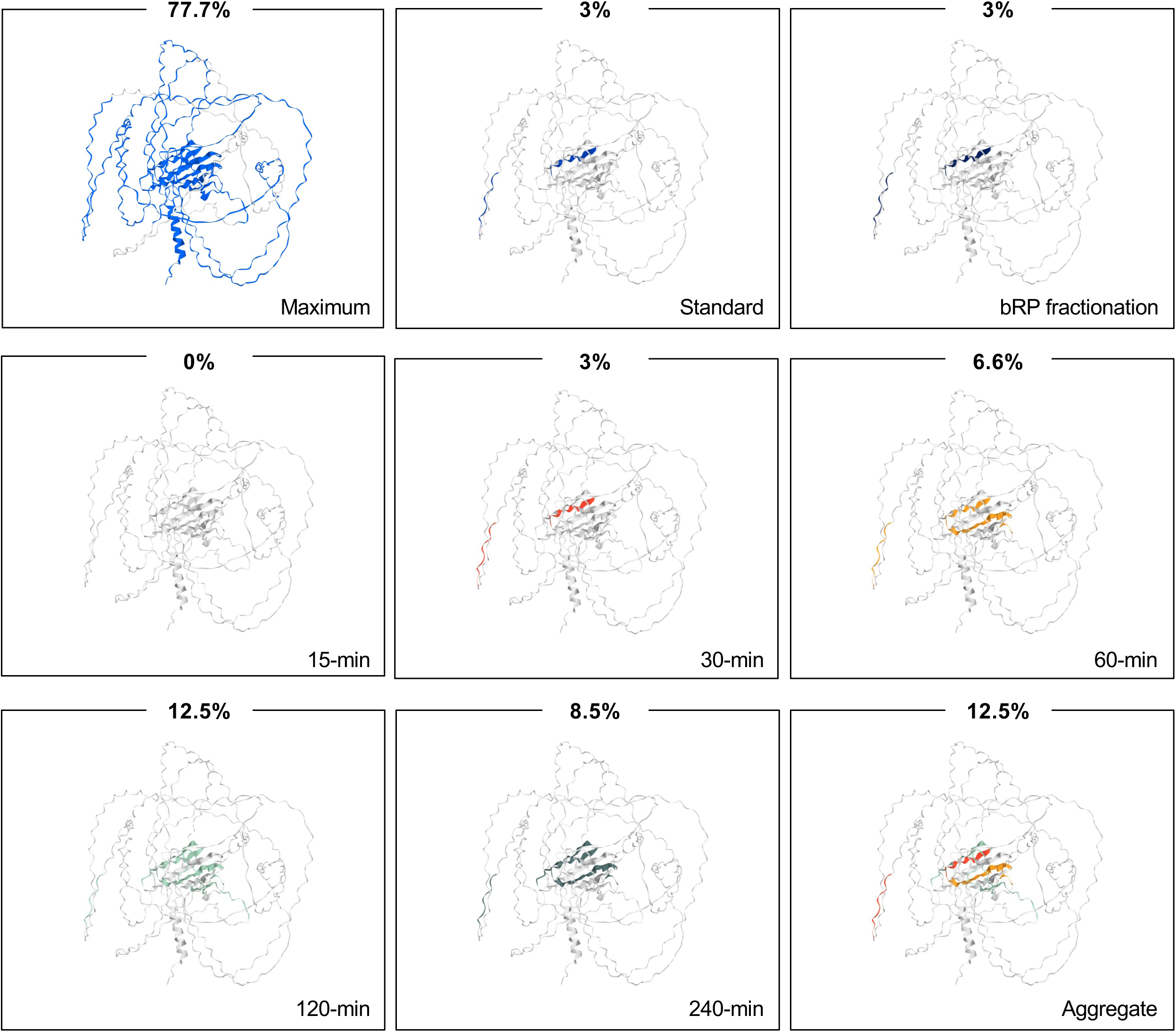
3D visualization of the sequence coverage of the alpha-1 chain of collagen VIII (*Col8a1*) achieved using different experimental pipelines. Panels represent the overlay of peptides detected using the standard tryptic digestion pipeline, the bRP fractionation pipeline, or at different timepoints (15 min: dark red, 30 min: light red, 60 min: gold, 120 min: teal, and 240 min: dark green) of the time-lapse tryptic digestion pipeline, onto the AlphaFold-predicted structure of the alpha-1 chain of collagen VIII (Col8a1) (UniProt Q00780; https://alphafold.ebi.ac.uk/entry/Q00780). The average sequence coverage calculated is indicated for each experimental condition. The “maximum” panel depicts the theoretical sequence coverage predicted via *in-silico* digestion. Peptides are color-coded in the “Aggregate” panel based on the first timepoint at which they were detected.

**Figure 7.**
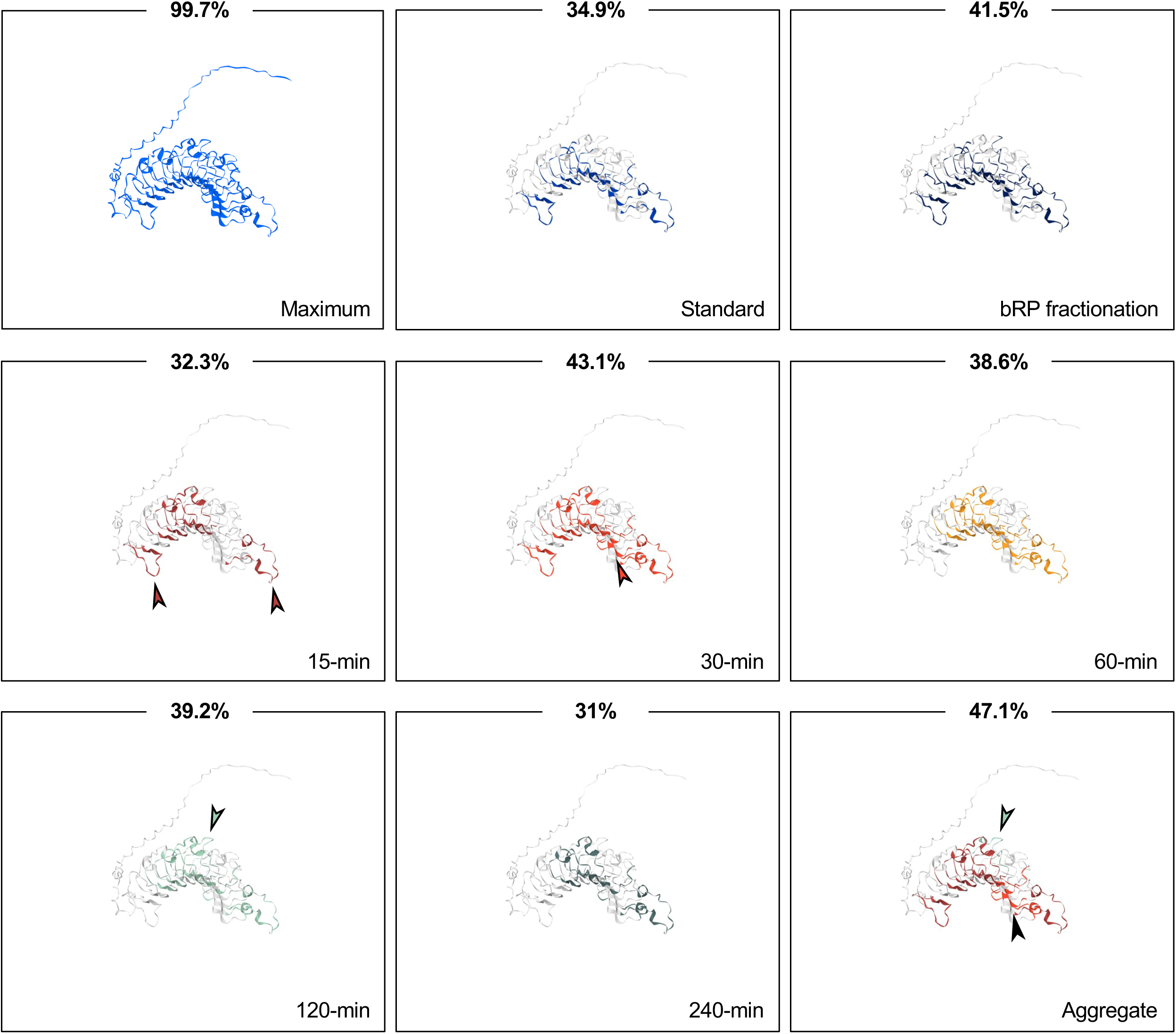
3D visualization of the sequence coverage of the proteoglycan prolargin achieved using different experimental pipelines. Panels represent the overlay of peptides detected using the standard tryptic digestion pipeline, the bRP fractionation pipeline, or at different timepoints (15 min: dark red, 30 min: light red, 60 min: gold, 120 min: teal, and 240 min: dark green) of the time-lapse tryptic digestion pipeline, onto the AlphaFold-predicted structure of prolargin (also known as proline-arginine-rich end leucine-rich repeat protein encoded by *Prelp*, UniProt Q9JK53; https://alphafold.ebi.ac.uk/entry/Q9JK53). The sequence coverage calculated is indicated for each experimental condition and is calculated as the proportion of the sequence covered by peptides detected in at least three of the five biological replicates. The “maximum” panel depicts the theoretical sequence coverage predicted via *in-silico* digestion. Peptides are color-coded in the “Aggregate” panel based on the first timepoint at which they were detected. Arrowheads point to peptides uniquely contributed by a given tryptic digestion timepoint.

**Figure 8.**
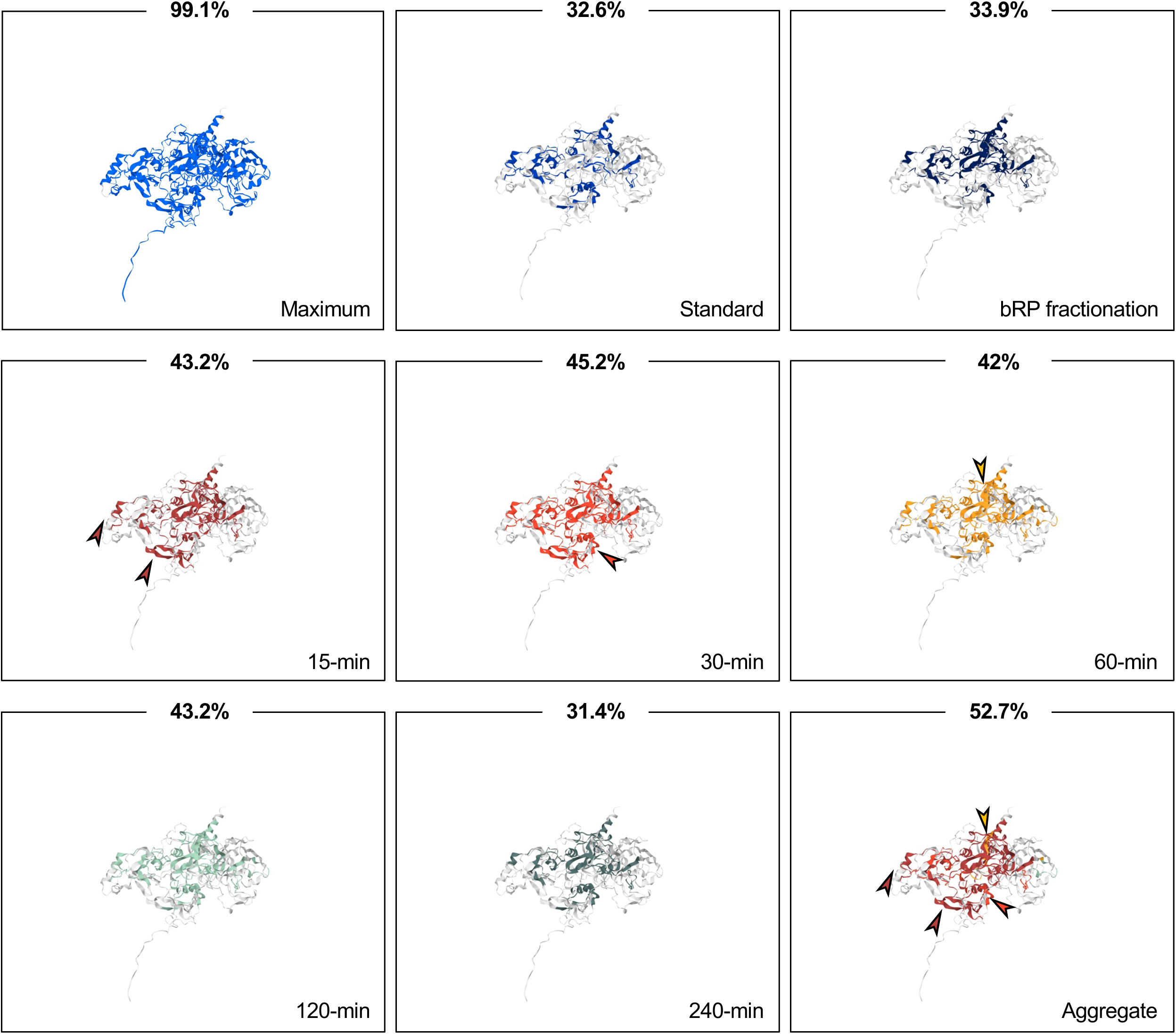
3D visualization of the sequence coverage of lysyl oxidase-like 4 achieved using different experimental pipelines. Panels represent the overlay of peptides detected using the standard tryptic digestion pipeline, the bRP fractionation pipeline, or at different timepoints (15 min: dark red, 30 min: light red, 60 min: gold, 120 min: teal, and 240 min: dark green) of the time-lapse tryptic digestion pipeline, onto the AlphaFold-predicted structure of lysyl oxidase-like 4 (UniProt Q924C6; https://alphafold.ebi.ac.uk/entry/Q924C6). The sequence coverage calculated is indicated for each experimental condition and is calculated as the proportion of the sequence covered by peptides detected in at least three of the five biological replicates. The “maximum” panel depicts the theoretical sequence coverage predicted via *in-silico* digestion. Peptides are color-coded in the “Aggregate” panel based on the first timepoint at which they were detected. Arrowheads point to peptides uniquely contributed by a given tryptic digestion timepoint.

The peptides used for 3D mapping correspond to those detected in at least three of the five biological replicates performed for each timepoint. Of note, the calculated percentage of sequence coverage indicated in **Figures 5 through 8** may thus slightly differ from the one obtained by simply averaging sequence coverage across biological replicates (**Supplementary Table 3**). Overall, the sequence coverage achieved by aggregating the peptides identified with the time-lapse digestion peptides (*see “Aggregate” panels*) are 59.9% for Postn, 52.7% for Loxl4, 47.1% for Prelp, but only 12.5% for Col8a1. These values are greater than those obtained using the standard bRP peptide fractionation protocol.

In addition to peptides detected in more than one timepoint of the time-lapse digestion, all examples demonstrate the unique contribution of different digestion timepoints to the release of different peptides and to increasing sequence coverage. In the case of periostin, a very short 15-minute incubation in trypsin is sufficient to achieve a 35.2% sequence coverage, and some of the peptides released at this early timepoint are not identified at the other timepoints (**Figure 5**, “15 min” panel, arrowhead). The maximum coverage is achieved at the 30-minute timepoint, but unique peptides are also released at the 2-h timepoint (**Figure 5**, “120 min” panel). Interestingly, the peptide uniquely contributed by the 15-minute timepoint maps to a protein loop that extends outside the core structure of the protein and, hence, is perhaps more readily accessible to tryptic digestion. For prolargin, the two earliest timepoints, 15 and 30 minutes, are sufficient to reach maximal experimental coverage since no unique peptide is contributed by the longer timepoints, as depicted on the “Aggregate” panel, where peptides are color-coded based on the first timepoint at which they were detected (**Figure 7**). Similarly, for Loxl4, the three earliest timepoints, 15, 30, and 60 minutes, are sufficient to reach maximal experimental coverage since no unique peptide is contributed by the 120-minute or 240-minute timepoints as illustrated on the color-coded “Aggregate” panel (**Figure 8**). Readers interested can retrieve peptide sequences of any of the proteins identified in this study (**Supplementary Tables 3D and 3E**) to perform similar 3D peptide mapping.

## DISCUSSION

In this study, we report the development of a time-lapse digestion protocol coupled with DIA-mass spectrometry to enhance ECM protein sequence coverage and gain insight into ECM protein folding in the context of an assembled ECM meshwork. We further highlight how time-lapsed proteomic data can be leveraged via 3-D peptide mapping to gain insights into the accessibility of different protein regions to trypsin and can thus help infer information on protein folding and, in the future, the identification of possible sites of protein-protein interactions.

While proteomics has proven critical to decipher the complexity of ECM protein compositions, proteoforms are the true effectors of molecular functions (14, 17, 31). Thus, it is of paramount importance to devise new experimental and computational approaches aimed at identifying and quantifying proteoforms, which will require achieving deeper sequence coverage (13).

Modulation of the duration of the trypsin digestion is only one of many modifiable parameters of a multi-step proteomic workflow. While we demonstrated its benefits in terms of sequence coverage, there is still a “dark side” to the matrisome. Indeed, across the 100+ matrisome proteins identified in this study, the average sequence coverage achieved is only 30.4% and a median coverage value of 28.5%. We were the first to propose and show that hard-to-digest ECM proteins would benefit from sequential multi-protease digestion (in our case, LysC + trypsin) (11), which is now broadly adopted and even commercially available. On the model of what has been developed to achieve higher coverage of the intracellular proteome and based on the findings reported here, we propose that a sequential (or simultaneous) digestion with *multiple* proteases, beyond the use of the LysC-trypsin pair, could achieve a deeper matrisome coverage as recently proposed (32). However, this poses challenges, including optimizing reaction conditions to ensure protease compatibility and downstream computational analysis to support multiplexed digestion for accurate peptide identification (33). To tackle this limitation, top-down proteomics is a relatively novel MS-based modality based on the ionization of intact proteins instead of digested peptides, using electrospray ionization and the analysis of the protein fragmentation pattern to get better insights into proteoforms (34–36). However, most ECM proteins are too large and difficult to ionize and, hence, are not good candidates for top-down proteomics. Thus, to date, bottom-up proteomics is still the most effective method for ECM profiling.

Protein structure prediction has made tremendous progress with the emergence of computational tools and methods such as AlphaFold (29, 37) and RoseTTAFold (38). However, while these machine learning models work very well on soluble proteins, they have limited success with predicting the structure of ECM proteins, particularly in those with long disordered regions and especially in their native state within the insoluble ECM meshwork. Of note, none of the structures used in this study for peptide mapping are experimental structures. The paucity of experimental data on the 3D matrisome protein structures is primarily due to the large size and insolubility of these molecules. Interrogation of the protein data bank (PDB), the repository of three-dimensional structures of biomolecules (39), revealed that the structure of only 67 full-length matrisome protein monomers and 41 full-length matrisome protein multimers have been experimentally solved, with only eight being structures of core structural matrisome components. Importantly, these experimental 3D structures have been acquired on the soluble form of the proteins, which are likely to adopt conformations vastly different from the conformations adopted in the context of the native insoluble ECM scaffold. The limitations of working with predicted structures for peptide mapping are particularly pronounced for multimeric proteins, such as triple-helical collagens (40). For example, the AlphaFold-predicted model of Col8a1 used for this analysis is a monomer, which exists only transiently *in vivo* when collagen chains are translated and is never found, as such, in the insoluble ECM. We thus propose that our approach can be used to refine predicted protein structures by providing additional restrictions to the prediction models; in turn, predicted structures can be validated with our experimental data using our time-lapse digestion data points and simple trypsin digestion models.

The complex structure of the ECM meshwork, established through protein-protein interactions, is critical to its function. Conventional structural proteomic approaches such as cross-linking mass spectrometry (41, 42) are unsuitable for studying ECM as cross-linking reagents will further challenge already hard-to-digest ECM proteins. On the other hand, the use of classical limited-proteolysis-coupled to mass spectrometry method developed to elucidate protein structures, conformational changes, and map sites of protein-protein interactions, has been limited to the study of soluble proteins as they provide omnidirectional access for proteases (43, 44). While the time-lapse digestion approach developed here can shed light on proteins’ accessibility in the assembled state of the ECM, as inferred by the proteins’ digestibility by trypsin, it is worth noting that it is applied here to samples that have been partially denatured. In the future, it will become important to devise methods to work on native ECM samples to conduct a true topographical survey of the ECM using proteomics. In addition, in native conditions, protein domains engaged in protein-protein interactions will be inaccessible and, as a result, resistant to proteolysis. Peptide mapping should thus reveal, more directly, a map of potential protein-protein interaction sites at the ECM scale.

Looking forward, we propose that applying time-lapsed proteolysis coupled with mass spectrometry can contribute to unraveling the compositional and structural complexity of the ECM with a depth never achieved before. This is a necessary first step toward deciphering ECM proteoform-dependent pathophysiological processes and the development of therapeutic strategies aiming at modulating ECM protein conformations, interactions, and signaling properties.

## EXPERIMENTAL PROCEDURES

### Cell culture and generation of cell-derived matrices (CDMs)

Previously characterized immortalized mouse embryonic fibroblasts (MEFs) isolated from our *Sned1^Neo-LacZ/Neo-LacZ^* mouse model and overexpressing the green fluorescence protein (45) were cultured in Dulbecco’s Modified Eagle Medium (DMEM, Corning, #10-017-CV) supplemented with 10 % fetal bovine serum (Sigma, #F0926), and 2 mM L-glutamine (Corning, #25-005-CI). Note that the genetic manipulation does not impair ECM production or assembly.

For each experimental condition, 10^6^ cells were seeded on a 6-cm dish and, upon reaching confluency (typically within 36 hours), were treated with 0.05 μg/mL ascorbic acid. Subsequently, half of the culture medium was replaced with fresh culture medium supplemented with 0.1 μg/mL ascorbic acid every other day for 7 days. On day 8, the cell cultures were decellularized as previously described (46). In brief, the cell layer was washed twice with phosphate buffer saline (PBS), then incubated with 0.5 mL of extraction buffer (0.5 % Triton X-100, 20 mM NH_4_OH in PBS) for 7 minutes at 37°C. The extent of the decellularization was visually assessed using phase-contrast microscopy. When deemed complete, 0.5 mL of cold PBS was added. Cell-derived matrices (CDMs) obtained were collected by gently tilting the plate on ice and transferred into a conical tube containing 10 mL of PBS for washing. The first wash was performed at 4°C for 4 hours, then PBS was replaced after centrifugation at 4,000 rpm for 8 minutes at 4°C. This was repeated once after 4 hours and then after 12 hours. After the washes, CDMs were pelleted in a 1.5-mL microcentrifuge tube by centrifugation at 13,000 rpm for 2 minutes at 4°C and stored at - 80°C until further processing.

### Sample preparation for mass spectrometry analysis

CDMs (n=5; replicates A to E) were resuspended in 100mM NH_4_HCO_3_ containing 8M urea and 10mM dithiothreitol (Thermo Scientific, #A39255) and incubated at 37°C for 2 hours under agitation as previously described (47). Partially solubilized CDMs were then alkylated by the addition of 25mM iodoacetamide (Thermo Scientific, #A39271) and incubated at room temperature for 30 minutes in the dark. Urea concentration was brought down to 2M with the addition of 100mM NH_4_HCO_3_ and samples were deglycosylated with PNGase F (New England Biolabs, #P0704L) for 2 hours at 37°C under agitation.

#### Standard tryptic digestion

Partially solubilized and deglycosylated CDMs were digested with trypsin (Thermo Scientific, #90058) overnight at 37°C under agitation (n=5) as previously described (47). Samples were centrifuged at 13,000 rpm for 5 minutes to separate any remaining undigested CDM material and peptide-containing supernatants were transferred to a clean tube. Samples were then acidified with 50% trifluoroacetic acid (TFA) and stored at −20°C until further processing.

#### Time-lapse tryptic digestion

For the time-lapse tryptic digestion protocol, CDMs were digested with trypsin (n=5) as follows: trypsin was added to each sample, and samples were incubated at 37°C under agitation. After 15 minutes, samples were centrifuged at 13,000 rpm for 5 minutes, and the supernatants containing tryptic peptides released were transferred to a clean tube, while the remaining partially proteins were resuspended in 2M urea in 100 mM NH_4_HCO_3_, with the addition of an equal amount of trypsin. After 15 minutes, the process was repeated to collect the 30-minute samples, and the same process was repeated to collect the 60-minute, 120-minute, and 240-minute samples. Peptide samples collected at each timepoint were acidified with 50% TFA.

#### High-pH reversed-phase peptide fractionation

For each sample digested with the standard protocol (n=5), 100 µg of peptides were further fractionated into 10 fractions eluted with the following concentration of acetonitrile in 0.1 % triethylamine: fraction 1: 5%, fraction 2; 7%, fraction 3: 9%, fraction 4: 11%, fraction 5: 13%, fraction 6: 15%, fraction 7: 17%, fraction 8: 20%, fraction 9: 25%, fraction 10; 50 % using a high-pH reversed-phase peptide fractionation kit (Pierce, #84868). The resulting fractions were then paired in a concatenated fashion (*i.e.*, fractions 1+6, fractions 2+7, fractions 3+8, fractions 4+9, and fractions 5+10).

#### Peptide desalting *and* concentration measurement

Acidified peptides were desalted using peptide desalting spin columns according to the manufacturer’s instructions (Pierce, #89852) and eluted in a 50% acetonitrile (ACN) solution containing 0.1% TFA. Peptides were lyophilized for 4 hours and reconstituted in a solution of 5% ACN and 0.1% formic acid. Peptide concentration was measured using a colorimetric assay (Pierce, #23275).

### LC-MS/MS analysis

#### Data acquisition

500ng-equivalent of peptides from each sample was dissolved in 10µL LC-MS grade 0.1% formic acid, injected, and separated by a capillary C18 reverse-phase column of the mass-spectrometer-linked Ultimate 3000 HPLC system (Thermo Fisher) using a 60-minute gradient (5 % - 85 % acetonitrile in LC-MS grade water) at a 300nL.min^-1^ flow rate. These samples were subjected to electrospray ionization (ESI) into a Q Exactive HF Orbitrap mass spectrometry system using data-dependent (DDA) or data-independent acquisition (DIA). Instrument control and data acquisition were performed via Xcalibur. DIA method consisted of a survey scan (MS1) at 45,000 resolution (automatic gain control 5e6 and maximum injection time of 200ms) from 350 to 1650 m/z followed by tandem MS/MS scans (MS2) with isolation window at 25 m/z. MS/MS scans were acquired at 30,000 resolution (automatic gain control target 5e5 and auto for injection time). The spectra were recorded in profile data type.

#### Predicted DIA-NN spectral library generation and file search

The *in-silico* predicted spectral library used in DIA analysis was generated by DIA-NN (version 1.8.1) (23), a deep-learning-based spectra and retention time prediction based on a custom proteome reference database generated by amending the standard mouse reference proteome (UniProt reference: UP000000589) with the GFP sequence. Trypsin with one maximum missed cleavage was enabled as *in-silico* cleavage rule, the maximum number of variable modifications was set to three, N-term M excision and C carbamidomethylation were enabled with Ox(M) and Ac(N-term) set as variable modifications. Peptide length was set within a range between 7-30 amino acids. The final spectral library (termed “DIA-NN” library in this manuscript) contained 55,381 protein isoforms and 7,670,070 precursors in 2,571,625 elution groups.

Mass spectrometry raw files were converted into .mzML files using msConvert (48) and searched using DIA-NN (version 1.8.1; see above) (23). Search settings were set as follows: Library generation was set to ‘smart profiling’, quantification strategy was set to ‘Robust LC (high precision)’, mass accuracies = 0.0 (automatic mass tolerances determination), Protein inference was set to ‘Genes’. ‘MBR’, ‘No shared spectra’, and ‘Heuristic protein inference’ were enabled.

In addition, precursor FDR was set as 1%. All other parameters were set as default. The report.*matrix result files containing identification and quantification matrices at the precursor, protein, and gene levels (FDR filtered at 1%) were subjected to downstream data processing and analysis.

#### Custom spectral library generation and file search

A custom spectral library (termed “DDA library” in the manuscript) was created by including raw files from a similar series of samples (*i.e.*, a dataset on the matrisome of MEFs generated using a standard and a time-lapse digestion protocol) acquired using a data-dependent mode (49). Datasets generated in this study were searched against this library using MSFragger (50), with fixed modification of carbamidomethyl cysteine and several variable PTMs enabled, including methionine oxidation, asparagine and glutamine deamidation, glutamine to pyroglutamate, and lysine and proline hydroxylation the latter two being characteristics of ECM proteins (13, 24). Trypsin was specified as the cleavage enzyme, allowing one missed cleavage; mass tolerance was set to 20 ppm for both precursor and fragment ions. After database search, a spectral library was generated using the in-house “Spec Lib” function in FragPipe (a GUI powered by MSFragger) with default settings. DIA raw files were searched against this custom-built spectral library, and search results at the peptide level were compared with those from searching against the predicted spectral library, in which no other variable PTMs were included other than methionine oxidation and N-terminal acetylation. Spectra of ECM peptides uniquely identified in the predicted DIA-NN spectral library were extracted from the predicted spectral library .tsv file and appended to assemble a larger custom-built spectral library supplemented with a set of raw files from a previously published study on the comprehensive proteomic characterization of the lung ECM (51) (see file list in **Supplementary Table 2**). This custom spectral library is termed “custom matrisome library” in the manuscript.

#### Data analysis

All analyses were restricted to reviewed UniProt entries and to proteins identified with two or more peptides. The dataset was annotated to identify ECM and ECM-associated proteins using the matrisome nomenclature (11, 52). Stringent criteria were selected for protein identification: the lists of proteins were aggregated within each of the five replicates by combining all proteins identified from each timepoint. These lists were then compared and the aggregate list of proteins from time-lapse digestion was defined as those identified in three of the five biological replicates.

##### Evaluation of protein abundance

Normalized total precursor ion intensities were used to evaluate protein abundance. The aggregated abundance of proteins was calculated within each replicate, where the aggregate was defined as the sum of the signals measured at each said timepoint (*e.g.*, aggregated relative abundance at 60 minutes was calculated by adding signal intensities of the 15-min, 30-min, and 60-min data points), with the values from five replicates and across the timepoints. Student’s t-test (two-tailed, unequal variance) was performed to determine statistical significance.

##### Analysis of sequence coverage

To investigate the aggregated coverage at a given timepoint of the time-lapse tryptic digestion, identified peptides from previous and current timepoints were pooled together prior to peptide mapping and coverage calculation (*e.g.*, aggregated coverage at 240 minutes was calculated based on all identified peptides from the sequential 15-min, 30-min, 60-min, 120-min, and 240-min digestions). Change in sequence coverage when comparing the standard tryptic digestion method and the data aggregated from the time-lapse tryptic digestion method was calculated as follows:

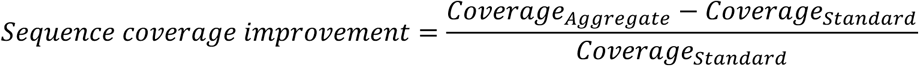

##### Statistical analysis

Considering the non-normality of the data, the distribution of sequence coverage values obtained with the standard 18-hour digestion or upon aggregation of the data from the time-lapse digestion pipeline were compared using the Wilcoxon signed-rank test. The coverage values of individual proteins were also compared using Multiple Mann-Whitney tests, with p-values adjusted with the Bonferroni-Dunn method for stringent results.

##### Peptide mapping on the 3D representation of protein structures

Number of peptides identified, the percentage of sequence coverage and peptide sequences are provided in **Supplementary Table S3**. Identified peptide sequences (**Supplementary Tables S3D and S3E**) were visualized and annotated using the Sequence Coverage Visualizer: https://scv.lab.gy/ (30). Maximum coverage was illustrated by visualizing peptides generated from *in-silico* digestion using ProteaseGuru (53), allowing one missed cleavage and peptides between 750 Da and 4,000 Da in size for consistency with the experimental search.

##### Data visualization

Proportional Venn diagrams were generated using BioVenn: https://www.biovenn.nl/ (54). The intersection of multiple datasets was visualized using UpSetR and the online application: https://gehlenborglab.shinyapps.io/upsetr/ (55).

## Data availability

Raw mass spectrometry data have been deposited to the ProteomeXchange Consortium (56) via the PRIDE partner repository (57) with the dataset identifier PXD061280. A comprehensive meta-data file in the Sample and Data Relationship Format (58) was also deposited to facilitate the re-use of our dataset. The raw data will be made publicly available upon acceptance of the manuscript.

## CRediT AUTHOR STATEMENT

**Fred Lee:** Formal analysis; Investigation; Methodology; Visualization; Roles/Writing - original draft.

**Xinhao Shao:** Formal analysis; Investigation; Methodology; Software; Visualization; Roles/Writing - original draft.

**James M. Considine:** Formal analysis; Investigation; Methodology; Roles/Writing - original draft.

**Yu (Tom) Gao:** Conceptualization; Formal analysis, Funding acquisition, Investigation; Methodology; Resources; Software; Supervision; Validation; Roles/Writing - original draft.

**Alexandra Naba:** Conceptualization; Formal analysis; Funding acquisition, Investigation; Methodology; Project administration; Resources; Supervision; Validation; Roles/Writing - original draft.

## ACKNOWLEDGMENTS

The authors would like to thank Chang Liu from the Gao lab for technical assistance and all the members of the Naba and Gao labs for helpful discussions.

## FUNDING SOURCES

This work was supported in part by the National Human Genome Research Institute (NHGRI) of the National Institutes of Health and the National Institutes of Health Common Fund through the Office of Strategic Coordination/Office of the NIH Director [U01HG012680 to AN and YG], the National Cancer Institute [R21CA261642 to AN and YG], the National Institute of Arthritis and Musculoskeletal and Skin Diseases [R01AR074997 to AN], and by the University of Illinois Cancer Center through the UICC Pilot Project fund [award 2020-PP-07 to YG and AN].

## CONFLICT OF INTEREST

AN has a sponsored research agreement with Boehringer-Ingelheim for work unrelated to the research presented in this study and holds a consulting agreement with AbbVie. The other authors declare that they have no conflict of interest.

## SUPPLEMENTARY TABLES

**Supplementary Table S1. List of raw MS files from Schiller *et al.*, 2015 used to build a custom matrisome spectral library**

**Supplementary Table S2**. **Complete MS dataset**

**S2A.** List of files and corresponding experimental conditions

**S2B.** Total precursor ion intensity values obtained using the DIA-NN library search strategy; *related to Figures 2A and 2C, and Supplementary Figure S2*.

**S2C.** Total precursor ion intensity values obtained using the DDA library search strategy; *related to Figures 2A and 2C, and Supplementary Figure S2*.

**S2D.** Total precursor ion intensity values obtained using the custom matrisome library search strategy; *related to Figures 2A and 2C, and Supplementary Figure S2*.

**S2E.** List of matrisome proteins identified in at least three of the five biological replicates generated using the standard overnight tryptic digestion pipeline; *related to Figures 2B and 2E.*

**S2F.** List of matrisome proteins identified in at least three of the five biological replicates generated using the time-lapse tryptic digestion pipeline; *related to Figure 2D and 2E*.

**Supplementary Table S3. Sequence coverage data and peptide sequences**

**S3A.** Sequence coverage values (%) for matrisome proteins detected in samples generated through time-lapse tryptic digestion pipelines and obtained using the custom matrisome library search; *related to Figure 3A*.

**S3B.** Sequence coverage values (%) for matrisome proteins detected in samples generated through the standard or the aggregation of the data from each timepoint of the time-lapse tryptic digestion pipelines and obtained using the custom matrisome library search; *related to Figures 3B and 4A*.

**S3C.** Cumulative sequence coverage values (%) for matrisome proteins detected in samples fractionated using basic reversed-phase liquid chromatography or generated using the time-lapse tryptic digestion pipeline and obtained using the custom matrisome library search; *related to Figures 3C and 4B*.

**S3D.** Peptide sequences corresponding to matrisome proteins detected in samples generated through the standard or time-lapse tryptic digestion pipeline and obtained using the custom matrisome library search; *related to Figures 4 to 7 and Supplementary Figures S3 and S4*.

**S3E.** Peptide sequences corresponding to matrisome proteins detected in samples fractionated using basic reversed-phase liquid chromatography and obtained using the custom matrisome library search; *related to Figures 4 to 7 and Supplementary Figures S3 and S4*.

## SUPPLEMENTARY FIGURE LEGENDS

**Supplementary Figure S1.**
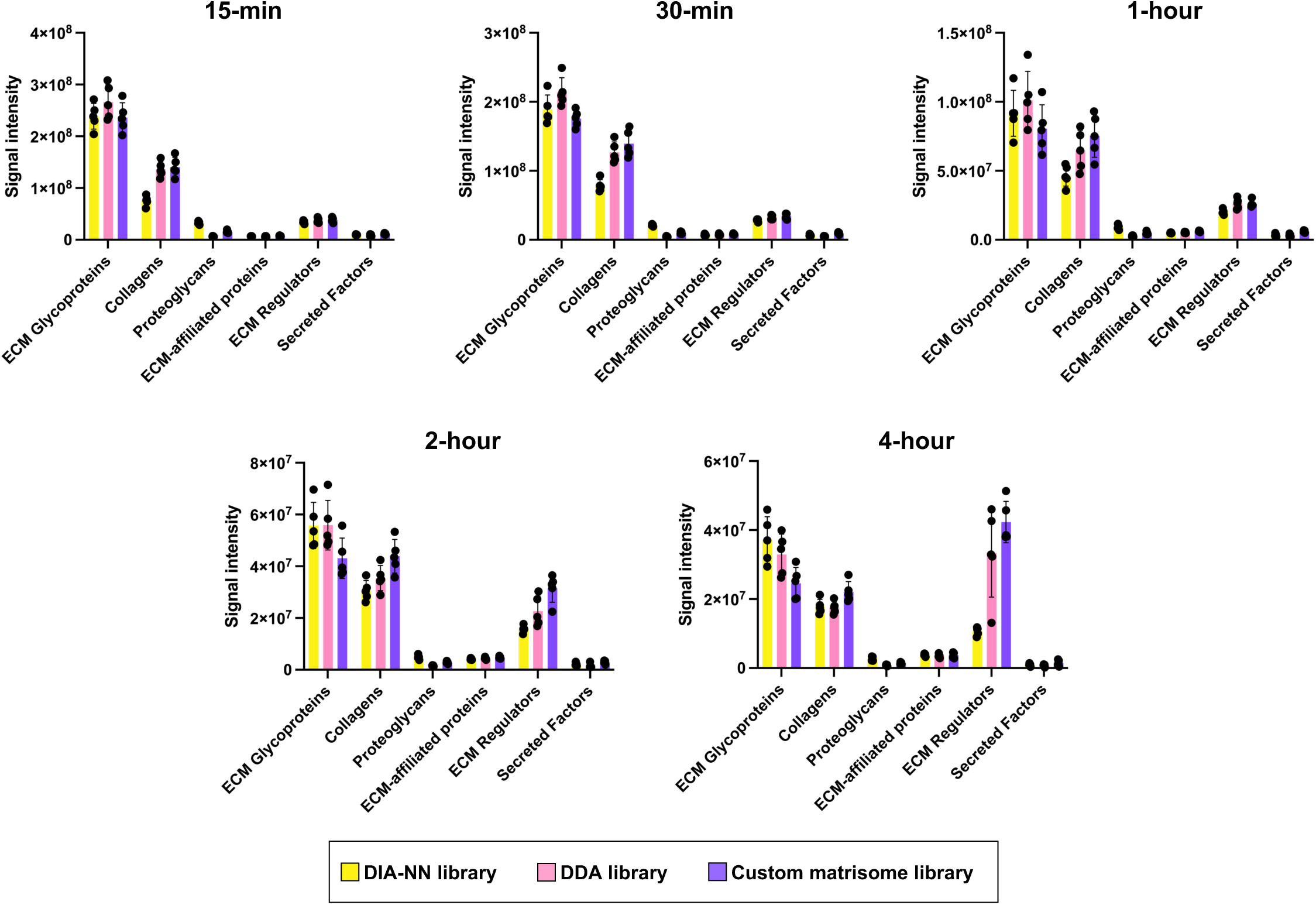
A custom spectral library enhances the identification of matrisome proteins in DIA datasets. Bar graphs represent the total precursor ion intensities measured by searching the datasets corresponding to the individual timepoints of the time-lapse tryptic digestion pipeline against each of the three spectral libraries assembled for this study and for each category of matrisome proteins.

**Supplementary Figure S2.**
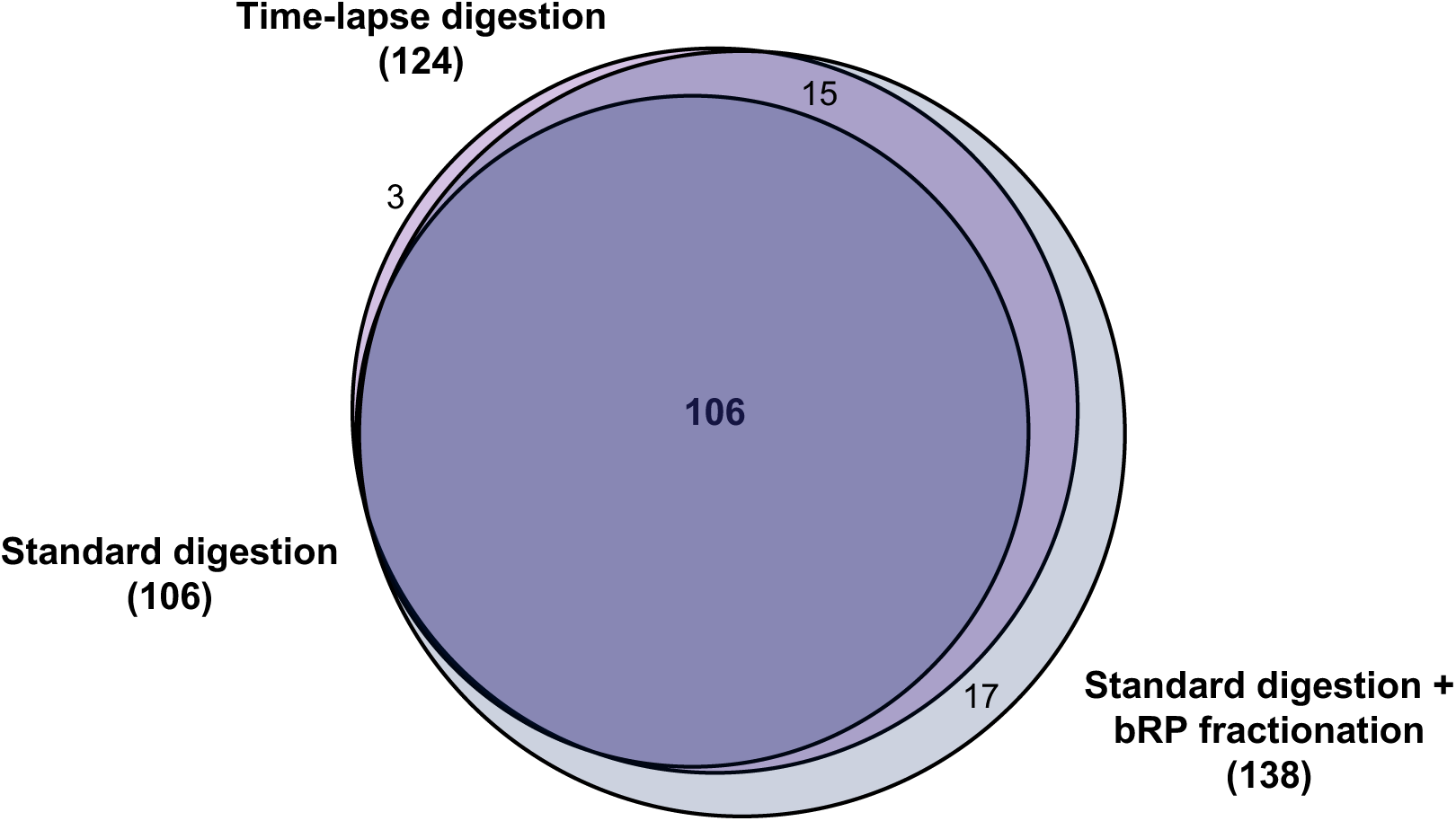
Comparison of the matrisome proteins identified using three different sample processing pipelines. Venn diagram represents the overlap between the matrisome proteins identified by searching the data generated through the standard overnight tryptic digestion pipeline (dark purple), the bRP fractionation pipeline (grey), or the aggregated data generated using the time-lapse tryptic digestion pipeline (little purple) against the custom matrisome spectral library. The lists of proteins composing each matrisome include proteins found in at least three of the five biological replicates.

## Notes

### Competing Interest Statement

The authors have declared no competing interest.

